# A LINK BETWEEN AGING AND PERSISTENCE

**DOI:** 10.1101/2024.08.24.609520

**Authors:** A. M. Proenca, C. U. Rang, L. Chao

## Abstract

Despite the various strategies that microorganisms have evolved to resist antibiotic treatments, most chronic infections are caused by subpopulations of susceptible bacteria in a transient state of dormancy. This phenotype, known as bacterial persistence, arises due to a natural and ubiquitous heterogeneity of growth states in bacterial populations. Nonetheless, the unifying mechanism of persistence remains unknown, with several pathways being able to trigger the phenotype. Here, we show that asymmetric damage partitioning, a form of cellular aging, produces the underlying phenotypic heterogeneity upon which persistence is triggered. Using single-cell microscopy and microfluidic devices, we demonstrate that deterministic asymmetry in exponential phase populations leads to a state of growth stability, which prevents the spontaneous formation of persisters. However, as populations approach stationary phase, aging bacteria — those inheriting more damage upon division — exhibit a sharper growth rate decline, increased probability of growth arrest, and higher persistence rates. These results indicate that persistence triggers are biased by bacterial asymmetry, thus acting upon the deterministic heterogeneity produced by cellular aging. This work suggests unifying mechanisms for persistence and offers new perspectives on the treatment of recalcitrant infections.

**IMPORTANCE:** Whenever bacterial cultures are treated with antibiotics, a fraction of the population survives despite exhibiting no active resistance mechanisms. These “persisters” are cells in a state of slow growth or dormancy, already present in the population prior to antibiotic exposure. Although various stressors or mutations increase persistence rates, a unifying persistence mechanism has not been established. Here, we show that cellular aging can represent such a mechanism. Bacteria age through the inheritance of intracellular damage, which occurs even in unstressed populations. As populations approach stationary phase, aging *Escherichia coli* have a steeper decline in elongation rates and earlier division arrest compared to younger cells. Upon antibiotic treatment, aging bacteria have higher persistence rates. These results show that stationary phase, a well-established persistence trigger, operates on the phenotypic heterogeneity produced by cellular aging. Because aging is a deterministic and ubiquitous process, it could represent a fundamental mechanism for the formation of persisters.

## INTRODUCTION

Antibiotic persistence is a ubiquitous phenotype in which subpopulations of susceptible bacteria survive drug treatments without being resistant. Formed due to phenotypic heterogeneity in bacterial populations, persisters are slow-growing or dormant cells present in the population prior to antibiotic exposure, generating an equally susceptible population after the treatment (*1–5*). While the immune system alone is usually able to eliminate persisters left behind by antibiotics, these bacteria represent a serious concern in immunocompromised patients or in infections that evade immunity (*6*). Nonetheless, despite the identification of a wide variety of persistence triggers and potential molecular pathways over the years, a unifying persistence mechanism has not been established.

Unlike the active evasion strategies employed by antibiotic resistant bacteria, persistence derives from the heterogeneity of growth states in bacterial populations (*7*). This distinct physiological state arises either spontaneously, as a stochastic phenotypic switch during steady-state growth (*8, 9*), or due to specific triggers, such as stationary phase (*2, 10, 11*), toxin production (*12, 13*), or damage accumulation (*14, 15*). Whereas the reports on spontaneous persisters are scarce and controversial (*5, 16*), persistence triggered by stationary phase has been well documented in *Escherichia coli* strains, particularly the high-persistence *hipA7* mutant (*2, 17, 18*). The accumulation of the HipA toxin in the *hipA7* mutant is reported to result in stochastic transitioning between dormant and growing states (*10*), with persister frequencies increasing in stationary phase. Similarly, a growth state transition can be induced by increased oxidative stress and protein misfolding (*14, 15*) in a process requiring the SOS response (*19*) and protein repair machinery (*9, 14, 20*). Recent advances indicate that ribosome dimerization plays a central role in this transition (*21, 22*), although the underlying source of heterogeneity on inactivated ribosomes remains puzzling. Overall, broad factors acting on bacterial heterogeneity could lead to the formation of persisters.

Contrary to the idea of stochastic switching between growth states, previous research suggests a clear deterministic correlation between non-genetic damage and growth physiology in rod-shaped bacteria (*23–25*). Misfolded proteins accumulate in the form of aggregates, which become anchored at the cell poles (*26*). Upon division, each cell inherits a newly synthesized pole, formed at the fission site, and an old pole carrying damage accumulated by the mother (Fig. 1A). On the following generation, one cell (*new daughter*) inherits the maternal new pole, carrying little damage, while its sibling (*old daughter*) inherits an old pole harboring larger damage loads. This asymmetric damage inheritance produces phenotypic heterogeneity, leading to deterministic aging and rejuvenation in bacterial populations (*25, 27, 28*). While in constant environmental conditions, old and new lineages reach distinct states of growth equilibrium (Fig. 1B) (*28, 29*). However, when a population is exposed to extreme levels of oxidative damage, cellular aging leads to a growth arrest in old daughters, whereas new daughters continue to replicate (Fig. 1C) (*30*). Although it has been suggested that this asymmetry could explain the formation of persisters (*31, 32*), few studies have attempted to trace such parallel.

**Fig. 1.**
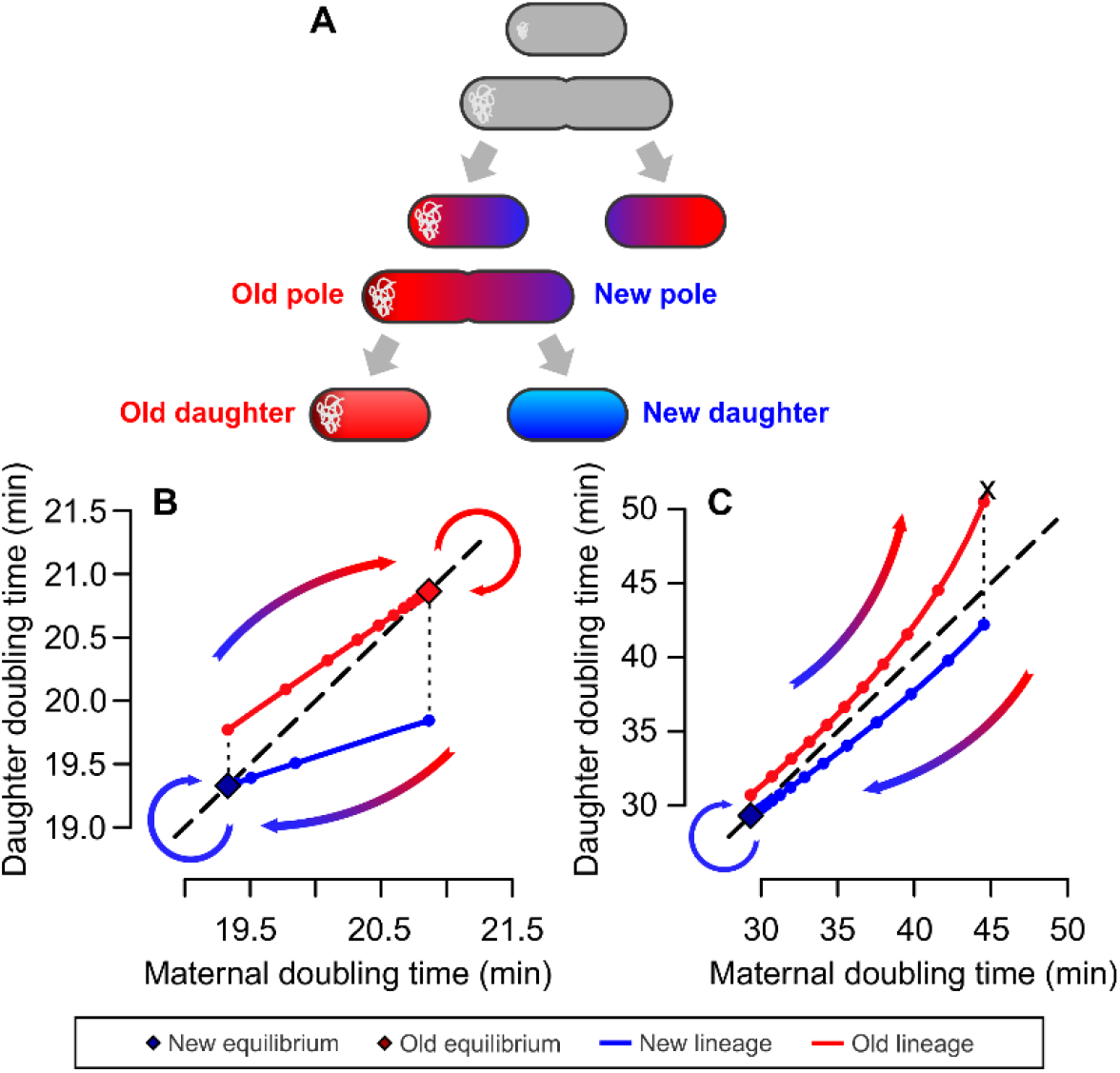
Cellular aging drives phenotypic transitions in bacteria. **(A)** Bacterial aging is a function of cell pole inheritance. Whenever a cell divides, it gives to its daughters a new pole, generated from the fission site, and an old pole that carries accumulated damage. The inheritance of either pole upon the next generation can be used to differentiate old daughters from new daughters. **(B and C)** Phase planes depicting the transition between growth states dictated by aging and rejuvenation, combining mathematical modeling (*29*) and empirical parameterization (*30*). Model details are provided in the Supplementary Text **(B)** Due to damage inheritance, old daughters take longer to divide. Cells consecutively inheriting either new or old poles over generations stabilize around new or old growth equilibria. Cells age or rejuvenate as they move between equilibria. **(C)** Under oxidative stress, damage accumulation leads old lineages toward increasingly long doubling times, until division arrest. Through rejuvenation, new daughters continue to proliferate under the same conditions.

Here, we propose a unifying explanation for persistence by correlating this phenotype with asymmetric damage inheritance. Through single-cell microscopy and microfluidics, we followed individual *E. coli* lineages through processes of continuous replication, growth arrest, and persistence. We show that old daughters, by inheriting larger damage loads, comprise a subpopulation of slow-growing cells with higher sensitivity to persistence triggering. These results suggest a simple underlying mechanism for the formation of persisters due to deterministic aspects of cellular aging.

## RESULTS

### Spontaneous persistence does not occur during stable exponential phase

We started by investigating the spontaneous formation of persisters during exponential growth. Because cellular aging modulates bacterial physiology and phenotypic heterogeneity, we asked whether it could determine which cells transition to a dormant state. Additionally, direct observations of stochastic growth arrest are scarce, rendering spontaneous persistence a controversial phenomenon. To address and control for these issues, we first followed wildtype MG1655 *E. coli* lineages in microfluidic devices. These devices capture bacterial cells while providing them with fresh culture medium throughout the experiments, allowing for long-term growth stability under single-cell microscopy. We combined data from two microfluidic designs: the mother machine (*33*), which traps old daughters at the bottom of growth wells, and the daughter device (*34*), with large chambers that contain 2D colonies trapping old and new daughters alike (Fig. S1). By following new and old daughters over generations, we looked for bacteria that stochastically switched between growing and dormant states.

The low frequency of spontaneous persisters poses a challenge for single-cell imaging and requires a substantial number of observed cells. Thus, we measured elongation rates and doubling times of 11,496 individual bacteria on 19 experimental replicates. Under continuous exponential growth, we observed stable elongation rates over generations (Fig. 2A). Old daughters displayed slower growth (0.978 ± 0.071 min^-1^) than new daughters (1.021 ± 0.064 min^-1^; one-tailed t test, t = 34.614, df = 11,177, p < 0.001), with an asymmetry that remained consistent throughout the observations. To better visualize this asymmetry, we represented the doubling times of each mother and its daughters on a phase plane (Fig. 2B), observing a separation between subpopulations of new and old daughters.

**Fig. 2.**
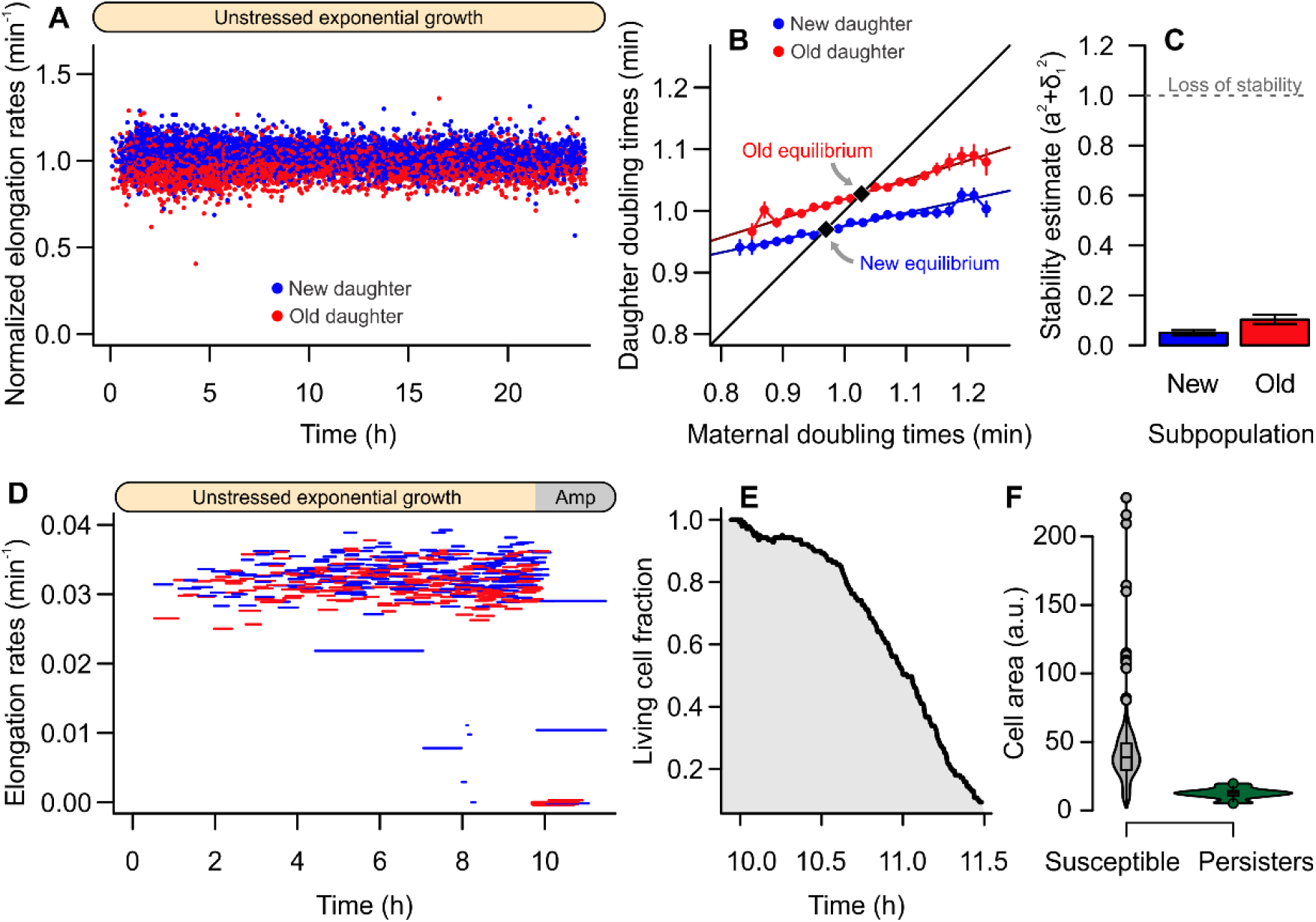
Absence of persistence during exponential phase. **(A)** Unstressed bacterial populations had stable elongation rates over time, without signs of transient growth arrest (n = 11,496 cells). Data were normalized to combine populations from distinct microfluidic designs. **(B)** Old daughters displayed longer doubling times (1.024 ± 0.078 min, n = 5,670) than new daughters (0.978 ± 0.064, n = 6,195; one-tailed t test, t = 34.505, df = 11,037, p < 0.001). These subpopulations reached distinct points of predicted physiological equilibrium, in which the doubling time of the mother equals that of a daughter. **(C)** Both new and old equilibria were stable, suggesting that these lineages are not subject to stochastic growth arrest (see Methods and Fig. S2 for details). **(D and E)** Once cells reached this state of stability, no persisters were observed upon exposure to 100 μg/ml Ampicillin. **(F)** On the other hand, when the antibiotic was introduced before cells reached stability, non-growing individuals (as indicated by cell area) carried over from the previous stationary phase persisted the treatment.

This representation allowed us to further investigate the physiological stability of these populations. New and old subpopulations are best described by separate regression lines, as shown in Fig. 2B and Fig. S2. When a regression with slope < 1 intersects the identity line, this intersect illustrates a predicted point of equilibrium. At this point, the doubling times of mother and daughter are equal. Lineages that continuously inherit either new or old poles should converge toward their respective equilibrium, remaining stable over time (*27, 28*). Although the constant elongation rates in Fig. 2A already suggested growth stability, spontaneous growth arrest could occur if stochasticity (*e*.*g*., of damage inheritance and partitioning) were sufficiently large. In our data, this would manifest as a loss of equilibrium, in which a given lineage loses stability and arrests division. Thus, to verify whether the equilibria predicted on Fig. 2B were stable, we estimated the stochasticity acting on both regression slopes (Fig. S2). A lineage remains in physiological equilibrium as long as the sums of squared slope (a) and stochasticity (σ_1_) is lower than 1 (see Methods for details). For exponentially growing bacteria, both new and old subpopulations satisfied the stability condition a^2^ + σ_1_^2^ < 1 (Fig. 2C, Fig. S2). Consequently, no stochastic switching between active and dormant states was present in this system.

Because high-persistence mutants are often used in batch culture experiments to increase persister yields, we repeated the above experiments with a MG1655 *E. coli hipA7* mutant. The *hipA7* mutation reduces binding of the toxin HipA to the antitoxin HipB, increasing the frequency of persisters by 1,000 fold (*35, 36*). We expected to find higher levels of stochasticity in its growth physiology, which would explain its behavior of stochastic switching between active and dormant states. However, we found that *hipA7* mutants are as stable as wild-type *E. coli* (see Supplemental Materials and Fig. S3). Following loading onto microfluidic devices, the only remarkable difference between strains was that *hipA7* mutants displayed a longer lag phase (Fig. S3G).

To corroborate the absence of the stochastic switching between active and dormant state and persisters in our system, we exposed wild-type populations to antibiotic treatment. As expected, exponential phase populations subjected to 100 μg/ml Ampicillin yielded no persisters (Fig. 2DE). This outcome could be avoided by exposing cells to Ampicillin 1 h after loading into the mother machine device, rather than allowing bacteria to reach steady-state growth. A recovery period was initiated after 5 h of antibiotic exposure, and persister cells were identified as those that had produced growing lineages after 18 h of recovery. This approach yielded a persister frequency of 1.197%. Comparing cell sizes before antibiotic exposure (Fig. 2F), we observed a significant difference between susceptible (39.802 ± 18.727 a.u.) and persister cells (12.816 ± 4.015 a.u.; two-sample t test: t = 22.371, df = 20.734, p < 0.001). Since *E. coli* cells are naturally smaller during lag phase, these persisters were likely formed during stationary phase and had not initiated growth at the time of antibiotic exposure.

These single-cell observations suggest that, once all individuals of a bacterial population reach stable exponential growth, no persisters are present in the population. Cell lineages that have reached a state of physiological equilibrium continue to replicate, without stochastically switching to a dormant state.

### Persistence triggering favors the old daughter

Since unstressed populations of both wildtype and *hiA7* mutants in exponential phase did not exhibit enough phenotypic heterogeneity to generate persisters, we hypothesized that increasing the variability of growth states could aid in producing the phenotype. Previous studies have reported higher persister frequencies for batch cultures exposed to oxidative stress (*14, 15*), which could facilitate the observation of persistence under single-cell microscopy. Within the context of bacterial aging, exposure to oxidative damage increases variability and could induce growth arrest among a subset of individuals (*29, 30*) — with a predicted bias towards old daughters, since they have a higher probability of losing growth stability (Fig. 1C).

To test this hypothesis, we tracked cell lineages growing exponentially in mother machine devices and exposed the cells to two sets of damaging treatments. In the first treatment, cells were exposed to photo-oxidative stress (FITC), known to increase the heterogeneity of growth states in a population (*30*), followed by a lethal antibiotic treatment. The second treatment consisted of sublethal levels of Streptomycin, which induces protein misfolding and asymmetric partitioning between old and new daughters (*23, 37*), followed by lethal levels of Nalidixic acid (for details, see Supplemental Information). Despite both treatments being able to produce differential growth arrest, neither yielded persisters (Figure 3AB).

**Fig. 3.**
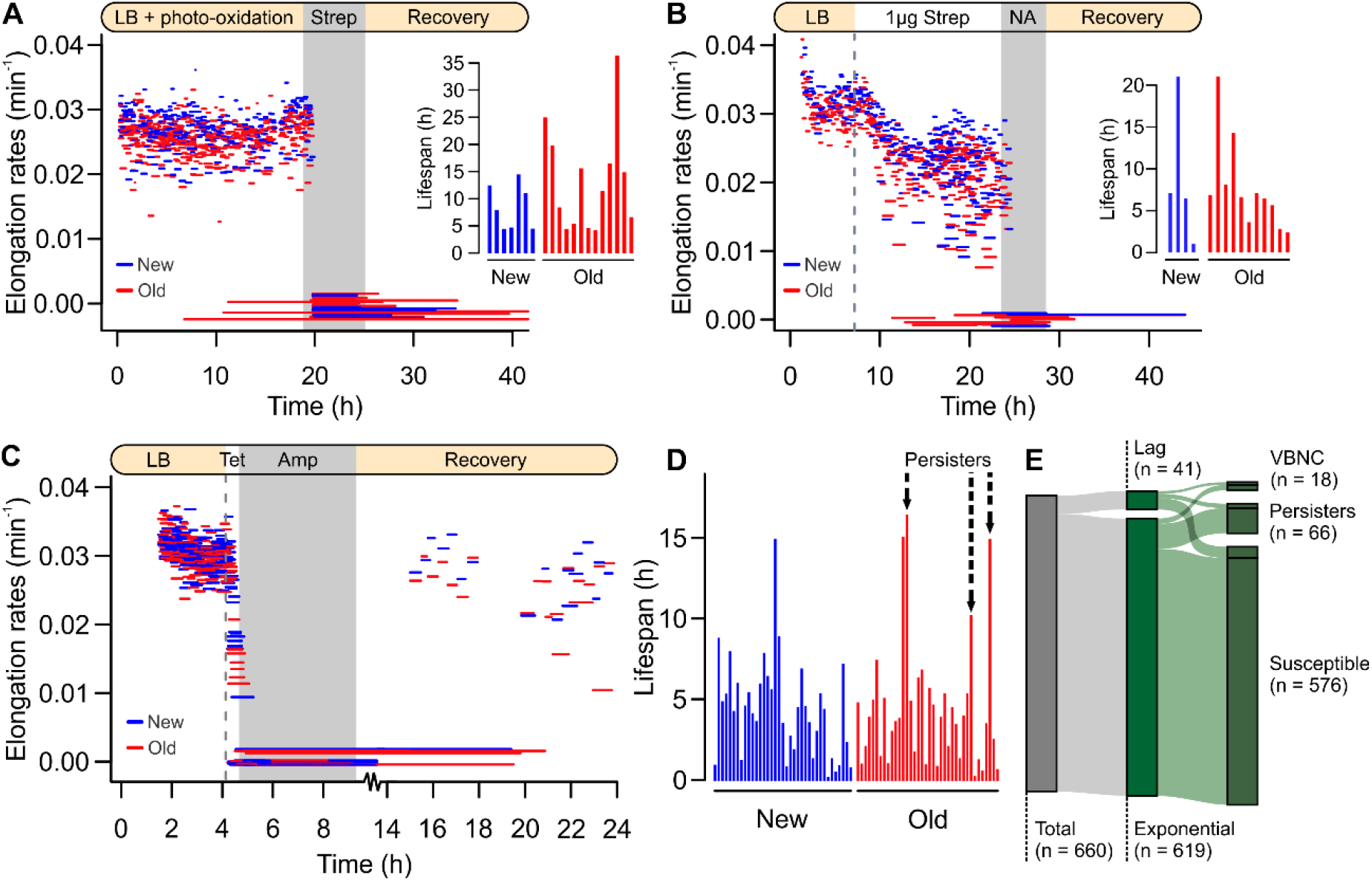
Old daughters favored by persistence triggering. Fig. 3 Triggered persistence from exponential phase populations. **(A)** Pre-treatment with photo-oxidative stress increased the difference between new and old daughter growth physiology (shown as elongation rates from birth to division or lysis). Upon exposure to 100 μg/ml Streptomycin and subsequent recovery, no persisters were observed. Detail: Lifespan of cells that arrested growth during treatment, until lysis. **(B)** Exposure to 1 μg/ml Streptomycin produced higher phenotypic heterogeneity (Fig. S4) prior to treatment with 30 μg/ml Nalidixic Acid (NA). No individuals resumed growth, suggesting that heterogeneity alone does not trigger persistence. **(C)** Using a known method to trigger persistence (*38*), we induced bacteriostasis through a short period of exposure to 50 μg/ml Tetracycline (dashed lines), followed by a treatment with 100 μg/ml Ampicillin (gray area). Whereas all cells arrested division through the bacteriostatic treatment, old daughter showed recovery once the treatment was removed. **(D)** Lifespan of cells that arrested growth, with persisters indicated by arrows. At the population level, the diagram **(E)** shows the total persistence triggered by Tetracycline followed by Ampicillin treatment.

Since enhancing growth heterogeneity and inducing damage accumulation failed to create persisters, we investigated whether a known persistence trigger could operate with a bias between old and new daughters. For this, we performed a pre-treatment with 50 μg/ml Tetracycline as described by Kwan et al. (*38*). The authors report that prior exposure to a bacteriostatic antibiotic increases persister formation frequency in exponential batch cultures, warranting its replication under single-cell microscopy. We followed the Tetracycline pre-treatment with 5 h of exposure to 100 μg/ml Ampicillin and observed whether bacteria resumed growth. Out of 28 lineages tracked through time-lapse microscopy (Fig. 3C), most cells lysed due to Ampicillin despite arresting growth during the Tetracycline pre-treatment. Nonetheless, once the antibiotic was removed, we observed persistence among old daughters (Fig. 3D). By further tracking 660 lineages through multi-field acquisition, we observed a population-wide persistence rate of 10% upon recovery (Fig. 3E). As in previous experiments, some individuals that arrested growth (2.73%) showed no recovery. Traditionally classified as viable but nonculturable (VBNC) cells, these most likely comprised dead individuals with an intact cell wall.

Taken together, these results suggest that inducing greater phenotypic heterogeneity is not enough to produce spontaneous persisters during exponential growth. A metabolic arrest trigger, such as Tetracycline, is essential to produce dormant cells that survive an antibiotic treatment. Interestingly, a persistence trigger might act based on the underlying phenotypic asymmetry, as indicated by the survival of old daughters. We further investigated this possibility in the following section.

### Stationary phase triggers persistence according to damage inheritance

Although the induction of bacteriostasis with Tetracycline was a successful persistence trigger, the narrow pre-treatment window (30 min of exposure) was not ideal for the study of cellular aging processes, which happen over generations. A triggering agent with chronic effects would be more suitable, as it would reflect natural processes of damage accumulation and asymmetric partitioning in a population. Stationary phase is a well-documented persistence trigger that satisfies this condition. As suggested on Fig. 2F and Fig. S3, stationary phase individuals are also a common confounding factor in persistence studies.

To test whether the underlying heterogeneity produced by cellular aging could interact with this trigger, we followed new and old daughters as they approached stationary phase. We replicated the transition from exponential growth to stationary phase by gradually diluting the culture media flowing through the microfluidic device, until it was nutrient-depleted (Fig. 4A) and cells entered stationary phase. These cells were then exposed to 100 μg/ml Ampicillin for 5.5 h. Through time-lapse microscopy, we tracked seven lineages that persisted through antibiotic exposure, with dormancy spanning between 6.8 and 8.5 h. Upon closer inspection of the transitioning phase between exponential and stationary phases (window detailed in Fig. 4B), we observed an interaction between time and age (new x old) as elongation rates decreased (one-way ANCOVA, F = 4.902, p = 0.0274). This suggests that the gradual growth arrest is best described by separate models, in which old daughters had a sharper decline (a = -0.0003, p < 0.001, R^2^ = 0.242) than new daughters (a = -0.0002, p < 0.001, R^2^ = 0.113).

**Fig. 4.**
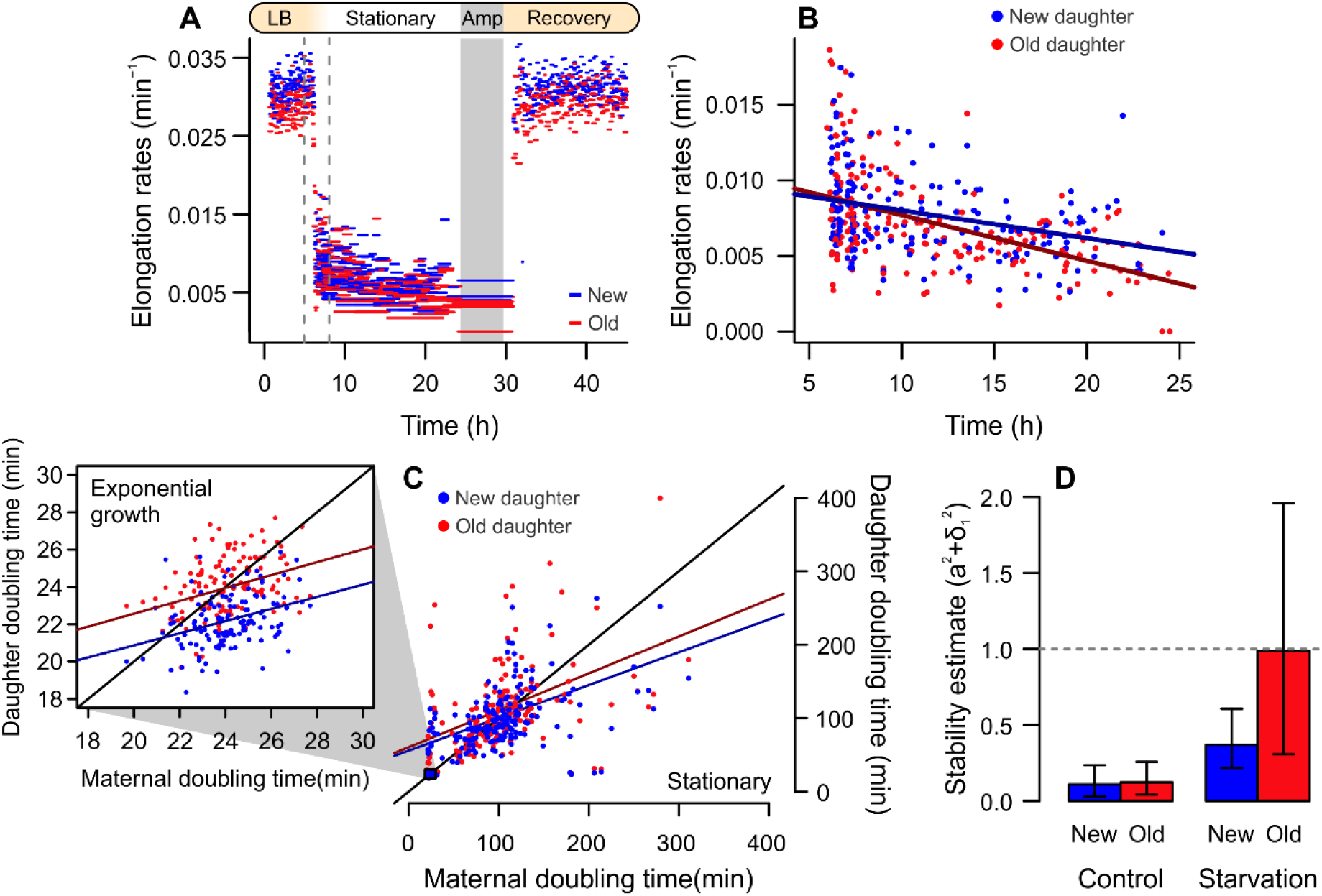
Old daughters are more sensitive to growth arrest triggered by stationary phase. **(A)** To induce a transition from exponential to stationary phase, cells were gradually starved of nutrients (dashed lines indicate transition interval), followed by Ampicillin exposure and recovery. **(B)** During starvation, old daughters continued to display slower elongation rates (0.0072 ± 0.0032 min^-1^, n = 196) than new daughters (0.0078 ± 0.0028 min^-1^, n = 183; one-tailed t test, t = 1.88, df = 374.34, p = 0.030). More importantly, old daughters displayed a steeper decline than new daughters, as indicated by linear models (solid lines). **(C)** There was an increase in doubling times and slopes between exponential (detail) and stationary phase (large phase plane). **(D)** As populations approached stationary phase, old lineages lost equilibrium (a = 0.502, σ_1_ = 0.857) and arrested growth, whereas new daughters continued to proliferate (a = 0.447, σ_1_ = 0.414). Error bars = 95% CI.

We analyzed the asymmetry of new and old subpopulations during this transition by contrasting their doubling times on phase planes (Fig. 4C). During exponential growth, the doubling time difference of 1.8 min between new (22.198 ± 1.379 min) and old daughters (23.973 ± 1.572 min) was significant (one-tailed paired t test, t = 12.474, df = 135, p < 0.001). This difference increased during starvation, with new daughters dividing every 102.234 ± 40.558 min and old daughters every 112.685 ± 53.097 min (one-tailed paired t test, t = 3.573, df = 167, p < 0.001). Because the linear regressions between maternal doubling times and daughter subpopulations displayed steeper slopes during starvation, we evaluated whether their equilibria lost stability (Fig. 4D). Whereas both new and old subpopulations exhibited growth stability during exponential phase, old daughters showed loss of equilibrium as they entered stationary phase (a^2^ + σ_1_^2^ = 0.988 [95% CI: 0.307-1.960]). A 10,000x bootstrapped analysis suggested that, due to combined effects of increased damage inheritance and stochasticity, old daughters had a 45.28% probability of arresting growth. New daughters had a higher probability of continuing to replicate (x^2^ = 5,850.6, p < 0.001), with an equilibrium that largely satisfied the stability requirement (a^2^ + σ_1_^2^ = 0.371 [95% CI: 0.217-0.606]). These results suggest that the transition to stationary phase triggers growth arrest with a bias towards old daughters.

To determine whether this growth arrest resulted in antibiotic persistence, we investigated the persistence rates of aging cells from stationary phase populations. Because stationary cells tend to move and rearrange within mother machine wells, tracking these lineages throughout antibiotic exposure and recovery is problematic. As an alternative approach, we let cells reach stationary populations in batch cultures. As a marker of cellular aging, we used cells expressing yellow fluorescent protein (YFP) bound to the small chaperone IbpA. This fluorescent reporter co-localizes with protein aggregates, indicating the presence of intracellular damage in each individual. Since large aggregates become anchored at old cell poles, IbpA-YFP acts as a marker for old daughters when there is enough damage accumulation in the system (*23, 26*). To induce non-lethal protein misfolding, we treated overnight cultures with 1 μg/ml Streptomycin (Fig. 5A), which increased the cell area occupied by aggregates from 4.27% to 10.34% (Fig. 5B; one-way ANOVA, F = 119.5, p < 0.001).

**Figure 5.**
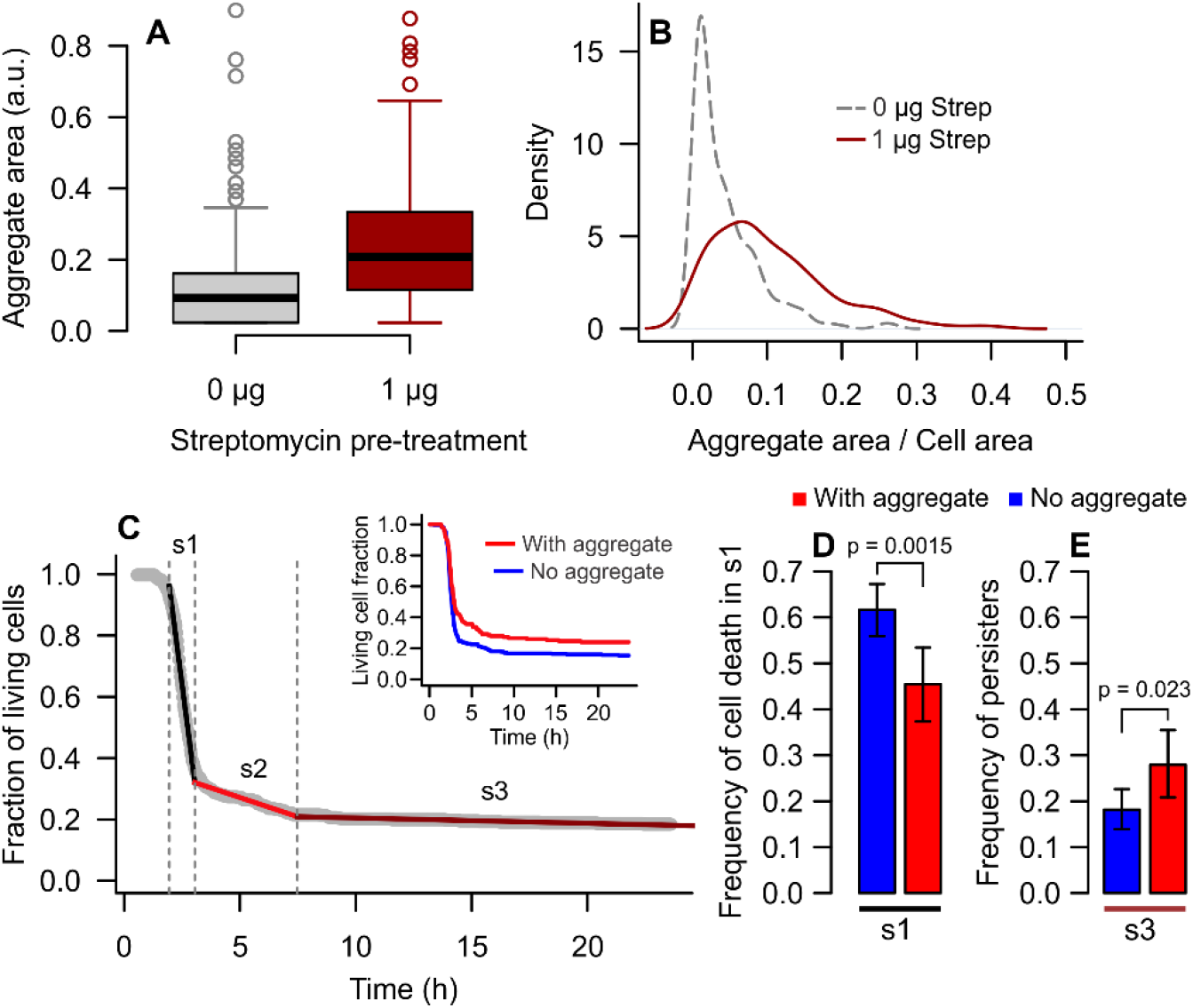
Stationary aging cells show higher persistence rates. **(A)** Damage accumulation was induced through overnight exposure to 1 μg/ml Streptomycin. Cells pre-treated exhibited larger aggregates (0.244 ± 0.171 a.u., n = 358) than non-treated cells (0.118 ± 0.126 a.u., n = 459; one-way ANOVA, F = 88.8, p < 0.001). **(B)** These aggregates occupied a larger area of the cell, thus increasing the anchoring of damage at old poles. This allowed for using IbpA-YFP-labeled aggregates as a marker for old daughters. **(C)** Cultures pre-treated with 1 μg/ml Streptomycin were loaded onto agarose pads containing 100 μg/ml Ampicillin and followed until lysis. A segmented fit to the killing curve indicated three break points (1.95, 3.05 and 7.48 h). Cells that survived to s3 were characterized as persisters. The detail shows separate killing curves according to the presence of aging markers. **(D)** The damage-free subpopulation had higher death rates from 1.95 to 3.05 h. **(E)** The frequency of persisters was higher among aging cells. Error bars = 95% CI.

These cells were inoculated onto agar pads containing 100 μg/ml Ampicillin and followed until lysis. Tracking cell counts over time, we observed a traditional killing curve (Fig. 5C) for which a segmented fit indicated three break points. A sharp population decline occurred from 2 to 3 h of exposure (s1 on Fig 5C) due to the quick lysis of susceptible cells. During this phase, aggregate-free individuals showed higher death rates (61.64%) than cells with aggregates (45.45%; Fig. 5D, x^2^ = 10.08, p = 0.0015). A slower population decline occurred from 3 to 7.5 h (s2 on Fig. 5C), followed by the stabilization of cell counts until the end of the experiment (s3 on Fig 5C). Cells that reached s3 were able to grow a new population when inoculated onto fresh LB medium, thus consisting of persisters. Splitting subpopulations according to damage loads, we observed that the subpopulation bearing aggregates stabilized at higher persistence rates (27.92%; detail in Fig. 5C, Fig. 5E) than the aggregate-free subpopulation (18.15%; x^2^ = 5.13, p = 0.023). These results suggest that individuals rejuvenated by asymmetry could become more susceptible to antibiotics, whereas aging cells have higher persistence rates.

Taken together, these results indicate that stationary phase triggers persistence with a bias produced by the asymmetric inheritance of damage. Deterministic asymmetry leads to heterogeneous bacterial populations, in which old daughters show slower elongation rates and earlier growth arrest entering stationary phase. Upon antibiotic treatment, this damaged subpopulation shows higher persistence rates. Therefore, bacterial aging offers a deterministic explanation for the underlying heterogeneity upon which persistence triggers promote antibiotic survival on subpopulations of bacteria.

## DISCUSSION

In this study, we established a link between antibiotic persistence and cellular aging. Both phenomena are ubiquitous among bacteria, deriving from the inherent phenotypic heterogeneity of unicellular populations. Here, we showed that the heterogeneity produced by the asymmetric partitioning of damage between new and old daughters provides the background upon which triggering agents induce persistence. Our microfluidic results indicate that unstressed, steady-state bacteria — whether bearing a high-persistence mutation or not — reach stable equilibria with distinct physiological states for new and old daughters (Fig. 2 and 3). However, in agreement with previous findings (*27, 28, 30, 33*), these populations retain long-term stability and proliferation, with no individuals exhibiting transient growth arrest. Although there are reports of spontaneous persisters forming during exponential phase in batch cultures, these rare observations are most likely due to the transfer of dormant cells from the previous stationary phase (*7, 16*). When these experiments are replicated in microfluidic devices, ensuring that all cells have reached exponential growth prior to antibiotic exposure, spontaneous persistence is not observed.

Through the investigation of stationary phase as a persistence trigger, we demonstrated that the asymmetry between new and old daughters leads to a differential response to starvation. Old daughters exhibited a sharper decline of elongation rates and entered a state of growth arrest, whereas new daughters continued to proliferate. In fact, previous studies have suggested that the old daughter phenotype accumulates as populations enter stationary phase (*31*), coinciding with an increase in persister frequencies due to the accumulation of intracellular damage (*14*). Moreover, while other studies have found that the deletion of DnaK chaperones decreases persistence rates (*9, 39*), we have previously observed that the same deletion causes mortality among old daughters (*30*). These prior observations motivated a closer inspection of deterministic aspects driving persistence patterns.

Although previous perceptions have favored stochasticity as the basis of the phenotypic heterogeneity leading to persistence (*6, 8, 40*), our results suggest that deterministic asymmetry represents the underlying persistence mechanism. This is not the first instance in which bacterial asymmetry arises as a deterministic explanation for the variability attributed to stochasticity. A similar shift in perspective occurred regarding the heterogeneity of elongation rates and protein synthesis in bacterial populations, which was formerly attributed to stochastic variation (*41, 42*). We have since consistently demonstrated that an essential component of this variance is produced by the inheritance and asymmetric partitioning of both damaged components (*28, 30, 43*) and newly synthesized proteins (*44*), thus providing a deterministic mechanism of phenotypic heterogeneity in clonal bacterial populations.

By transposing principles of cellular aging to the study of antibiotic persistence, bacterial aging itself acquires a new perspective. Bacteria were long regarded as immortal organisms, uncapable of aging. Through the observation of their divisional asymmetry, however, an old daughter becomes comparable to a disposable soma, which retains damage and favors de preservation of the germline (*45*). The maintenance of aging in complex organisms already poses an evolutionary puzzle, and even more so the prevalence of aging among unicellular organisms. Our work contributes to the elucidation of this phenomenon by showing that aging can be advantageous at the level of cell lineage survival. In bacteria, the retention of damage at the old poles is a cheap alternative to energy-intensive repair processes (*24*). Although this means that a portion of the population, the old daughters, has lower fitness, the resulting phenotypic heterogeneity has evolutionary advantages. *In silico* results have shown that asymmetric populations endure higher levels of stress than symmetric bacteria (*43*), and microfluidic experiments have demonstrated that lethal levels of oxidation lead to mortality in old lineages, while allowing for the survival of new daughters (*30*). Thus far, these results corroborated the parallels with the disposable soma hypothesis. Here, we showed another advantageous aspect of bacterial aging. As cells enter stationary phase, the bacteria that inherit more damage (non-lethal levels) arrest growth earlier and show higher persistence to drug treatments. Therefore, bacterial aging can lead to the differential survival of old daughters. This demonstrates that the deterministic heterogeneity produced by cellular aging is advantageous for bacterial populations.

The persistence of susceptible bacterial subpopulations to antibiotic treatments has long represented an elusive public health concern (*6, 46*). Through the elucidation of the underlying phenotypic heterogeneity through which extrinsic triggers lead to persistence, we thus propose cellular aging as a source of differential antibiotic survival. The application of this framework on future persistence research may contribute to the development of new strategies against recalcitrant infections.

## ACKNOWLEDGEMENTS

We thank A. Qiu, C. Shi and C. Buetz for data discussions and experimental assistance. We thank A. Lindner and K. Lewis for kindly providing strains. Work was supported by grants to L.C. from the National Science Foundation (DEB-1354253). This study was financed in part by the Coordenação de Aperfeiçoamento de Pessoal de Nível Superior - Brasil (CAPES) – Finance Code 001, through the Science Without Borders Fellowship (2014-2019) and the Program for Institutional Internationalization Postdoctoral Fellowship (2019-2021) provided to A.M.P.

## AUTHOR CONTRIBUTIONS

A.M.P, C.U.R., and L.C. conceptualized the work, designed methods and wrote the manuscript. A.M.P. and C.U.R performed experiments. A.M.P performed data collection, analysis, and visualization. L.C. provided resources and supervision.

## METHODS

### Bacterial strains and growth conditions

Experiments were performed using K-12 *E. coli* MG1655 and *E. coli* MG1655 *hipA7* (kindly provided by K. Lewis, Northeastern University, USA) as a high-persistence mutant. For the visualization of protein aggregates, we employed the strain MG1655 containing yellow fluorescent protein (YFP) bound to the small chaperone IbpA (*IbpA-yfp-Cm*^*r*^) (*23*). The construct was kindly provided by A. Lindner (INSERM, France), and inserted into the chromosome as previously described (*47, 48*). Bacteria were grown overnight in LB medium (lysogeny broth) at 37° C with agitation prior to loading into microfluidic devices or agar pads. For microfluidics experiments, LB media was supplemented with 0.075% Tween 20 to prevent the formation of biofilms on flow channels.

### Design and fabrication of microfluidic devices

Exponential growth experiments were performed using the mother machine or daughter device microfluidic designs (Fig. S1). The mother machine was based on the original design (*33*) and adapted by R. Johnson (Hasty Lab, UC San Diego). Each device consisted of 16 large flow channels containing 2,000 growth wells (1.25 × 30 × 1 μm) each. The daughter device was developed by O. Mondragón-Palomino (Hasty Lab, UC San Diego) (*34*) and contained 48 growth chambers (40 × 50 × 0.95 μm), each accommodating a monolayer of up to 300 cells at a time. Polydimethylsiloxane (PDMS; Kit Sylgard 184, VRW International, California) devices were fabricated through soft lithography from master silicon wafers used as negative molds. The mother machine silicon wafer was produced by Nano3 (UC San Diego), and the daughter device was provided by R. Johnson and the Hasty Lab (UC San Diego). PDMS devices had their loading ports punctured and were permanently attached to 24 x 40 mm coverslips through the exposure to O_2_ and UV light.

### Cell loading and damaging treatments

Overnight bacterial cultures were concentrated by centrifuging 1 ml of culture at 5,300 g for 2 min, followed by discarding of the supernatant media and resuspension into 50 μl LB-Tween 20 medium. The concentrated culture was loaded onto the device, ensuring that growth traps were filled and no air was left within the channels. For experiments with the *hipA7* strain, exposure to room temperature was minimized during loading due to cold sensitivity. After loading, inlet and outlet 60 ml syringes were connected to the ports, containing 30 ml of medium and 10 ml of MilliQ water, respectively. When necessary, the inlet was replenished or replaced throughout the experiments.

Exponential growth experiments were performed with continuous supply of culture media, starting with a 24 h control period that allowed all lineages to reach a stable growth state. Exponential persister assays followed this control period with a damaging pre-treatment, antibiotic exposure, and recovery in control conditions. Damaging pre-treatments consisted of sub-lethal stress exposure for 24 h, applied through fluorescent light excitation (FITC filter, 490 nm wavelength) in 2 min intervals, with exposure of 1.5 s, or 1 μg/ml Streptomycin added to the medium. The effect of these treatment on growth physiology has been described in detail (*30*). Alternatively, the pre-treatment was replaced by 50 μg/ml Tetracycline exposure for 30 min to induce growth arrest (*38*). Antibiotic treatments were performed with 100 μg/ml Streptomycin, 100 μg/ml Ampicillin or 30 μg/ml Nalidixic Acid for the indicated interval. Recovery periods represented a reversion to control conditions, thoroughly washing the inlet to remove traces of the antibiotic and providing fresh medium.

Stationary phase experiments in the mother machine device were achieved through a gradual dilution of LB medium with M9 minimal medium containing no glucose, over the course of 4 h. After 4 h, the inlet was washed with nutrient-depleted M9 to remove traces of LB, and cells were starved for the following 18 h. This treatment was followed by 100 μg/ml Ampicillin exposure for 5h and recovery under control conditions for the verification of persistence.

### Preparation and culturing on agarose pads

Stationary culture experiments presented in Supplementary Figure 3 were performed on LB agarose pads (LB medium supplemented with 15 g/l agarose). To induce protein aggregation (Figure 5), overnight cultures were grown in LB broth supplemented with 1 μg/ml Streptomycin, then loaded onto 10 μl agarose pads prepared immediately prior to the experiment. For persistence assays, the pad was supplemented with 100 μg/ml Ampicillin during this preparation. Cells were inoculated onto the agarose pads and followed over time until lysis. To ensure that the surviving cells were persisters, at the end of the experiment the agar pads were inoculated into LB medium and grown overnight.

### Time-lapse imaging

Cell movies were collected by a Nikon Eclipse Ti-S microscope, with imaging intervals controlled by NIS-Elements AR software. Mother machine and agarose pad phase images were obtained in 2 min intervals, while daughter device experiments were imaged every 20 s. When necessary for the visualization of protein aggregation, fluorescence images were obtained through a FITC filter (490 nm) following 20 min intervals to avoid damaging the cells.

### Quantification and data analysis

Images were analyzed using the software ImageJ (NIH, https://imagej.nih.gov/ij), recording cell coordinates as Regions of Interest (ROI) and cell names as indicatives of lineage and cell pole inheritance. Cell lengths were determined immediately before and after each division and time of division was recorded. Elongation rates (r) and doubling times (ln(2)/r) were calculated from the data, and the resulting tables were entered in an R program (version 3.4.1) (*49*) (R Core Team, https://www.r-project.org/) to determine maternity, sibling pairs and lineage trees. The ImageJ plugin MicrobeJ (*50*) (https://www.microbej.com/) was used for detection and segmentation of fluorescence images on the first image after loading onto agarose pads, before antibiotic exposure affected protein aggregation.

Data analysis was performed in R (version 3.4.1). Results are presented as mean ± standard deviation unless otherwise noted. Differences between new and old daughters were determined through one-tailed *t* tests, with paired data in the case of phase planes. New and old subpopulations of wild-type and *hipA7* strains were compared through two-way ANOVA. Confidence intervals (CI) obtained by bootstrapped analyses are presented for equilibrium stability estimates, variance partitioning and agarose pad persister frequencies. Elongation rates and doubling times in Fig. 2A and 2B were normalized to allow pooling of populations grown in either mother machines or daughter devices. The segmented fit and breakpoint estimation on Fig. 5C were performed using the “segmented” package in R (*51*).

### Equilibrium stability analysis

A detailed implementation of this analysis has been described in previous work (*28*). Briefly, when doubling times are presented on a phase plane, a separation between new and old subpopulations is observed whenever asymmetry is present. Phase planes are best described by linear regressions between the doubling times of the mothers and either new or old daughters. For regressions with a slope smaller than 1, an intersect with the identity line will occur. At this intersect, the doubling time of the mother equals that of a daughter inheriting the same pole. This point works as an *attractor* or *equilibrium point*, to which cells with stable growth return over generations. As long as the equilibrium is maintained, a cell lineage has immortal replication. Due to their physiological asymmetry, lineages consecutively inheriting either new or old poles reach distinct equilibria.

However, despite the deterministic effects of maternal damage inheritance and asymmetry, bacterial doubling times are highly stochastic. By chance, an individual in equilibrium might inherit a much larger damage load, with a physiological cost that will lead to a longer doubling time. In theory, if the stochasticity in the system is sufficiently large, it can lead to spontaneous events of growth arrest. This possibility must be considered when estimating the physiological stability of a lineage. Here, we define this factor as the stochasticity (σ_1_) acting on the slope (a) of each linear regression. To determine σ_1_, we compared slopes obtained for the entire subpopulations with the effective slopes connecting each mother (T0) and daughter (Ti) pair. This is given as (Ti – b)/T0 = a + ξ_1_, where ξ_1_ represents the deviation from the subpopulation slope. The standard deviation of all ξ_1_ values provides an estimate of σ_1_.

An equilibrium is stable provided that a^2^ + σ_1_^2^ < 1. If this condition is satisfied, daughters inheriting the same pole generation after generation tend to converge towards the equilibrium point. Otherwise, when a^2^ + σ_1_^2^ ≥ 1, stochasticity could lead the lineage to increasingly long doubling times, and eventual arrest of cell division. This process is graphically represented in Fig. 1B and 1C and has been experimentally verified (*28, 30*). Thus, the estimate of equilibrium stability for new and old subpopulations (as performed in Fig. 2C and 4D) provides a quantitative determination of whether lineages are able to sustain long term proliferation given the experimental conditions.

### Doubling time variance partitioning

The partitioning of doubling times variances into deterministic and stochastic components was performed according to Chao et al (*43*) and Proenca et al (*28*). While the former method calculates the absolute values of deterministic and stochastic components (Fig. S3), the latter estimates the relative contribution of these factors (Fig. S4).

For the variance partitioning presented in Fig. S3, the doubling time distributions of new and old daughters are normalized according to the doubling time of a hypothetical symmetric sibling, an intermediate between new and old subpopulations. With doubling times of the population centered around zero, the distance (*D*) between new and old doubling time distributions provides an estimate of the deterministic variance (*D*^2^/4). The mean variance of new (V_N_) and old (V_O_) doubling time distributions estimates the stochastic variance in the system ([V_N_ + V_O_]/2). This calculation allows for a comparison of populations of different sizes, but it does not take into account the maternal effect.

The variance partitioning method used for Fig. S4, on the other hand, uses a phase plane to extract deterministic and stochastic components, introducing the maternal doubling time as a deterministic factor. First, a single linear regression was traced between mothers and daughters of each population. The sums of squared deviations of predicted doubling times from the population mean estimates the maternal effect. Next, separate regressions were traced for new and old subpopulations, and the deviation of these predicted doubling times from the single population regression was estimated. A sum of these squared deviations represents the effect of asymmetry. Together, the maternal effect and asymmetry estimate the deterministic component. Finally, the stochastic component was estimated from the deviation of each experimental point from the doubling times predicted by new and old regressions. These values were expressed as fractions of the total variance, providing the relative contribution of each component.

## Data availability

All data supporting the conclusions of this study have been deposited on Dryad Digital Repository (doi: 10.5061/dryad.7d7wm3845) and will be made available by the time of publication.

