## Supplementary Materials for "A LINK BETWEEN AGING AND PERSISTENCE"

[2] Current address: Institute of Biology, Freie Universität Berlin; Berlin, 14195; Germany.

### SUPPLEMENTARY TEXT

#### Bacterial aging model

Aging and rejuvenation contribute to shaping the heterogeneity of growth states found within clonal bacterial populations (*27, 30*). To frame this age structuring, we can apply a population genetics model that describes bacterial growth and division across generations (*29*). For this, let us imagine that an *E. coli* cell is born free of intracellular damage ($k_0 = 0$). Over its lifetime, it accumulates damage at a rate λ:

$$k(t) = k_0 + \lambda t$$

This cell must also accumulate a certain amount of internal product in order to divide, similarly to an "adder" model (*52*). A damage-free cell would reach this checkpoint at time Π, the minimum possible doubling time ($T_0$):

$$\Pi = (1 - k_0)T_0 - (\lambda/2)T_0^2$$

Realistically, however, even cells cultured in benign environmental conditions accumulate intrinsic damage at a rate $\lambda > 0$ min$^{-1}$, which slows down the accumulation of internal product and leads to longer doubling times ($T_0$). Assuming that λ is linear, the cell will divide with a damage load $D_0$:

$$D_0 = k_0 + \lambda T_0$$

Upon division, this damage load is partitioned between the daughters with a certain degree of asymmetry (*a*). We define that *a* represents the fraction of $D_0$ inherited by the new daughter, such that $a = 0$ when the old daughter receives the full maternal load, and $a = 0.5$ for a symmetric division:

$$k_1 = D_0 a = (k_0 + \lambda T_0)a$$

$$k_2 = D_0(1 - a) = (k_0 + \lambda T_0)(1 - a)$$

The amount of damage inherited by new ($k_1$) and old ($k_2$) daughters, along with λ, will determine their respective doubling times ($T_1$ and $T_2$) at division:

$$T_i = \left\{(1 - k_i) - \sqrt{(1 - k_i)^2 - 2\Pi\lambda}\right\}/\lambda$$

The model thus allows us to estimate the doubling times of new and old daughters based on the initial damage load of the mother. To examine the core predictions generated from this mother, shown in Fig. 1, we used values for the parameters Π, λ, and *a* obtained from previous experiments (*30*). For an unstressed population, we assumed that Π = 18.5 min, λ = 0.0022 min$^{-1}$. For a population facing high levels of stress, we increased damage accumulation rates to λ = 0.0089 min$^{-1}$. Asymmetry values were assumed to vary with λ, as

$$a = 0.1007\ln(\lambda) + 0.9531$$

according to experimental observations (*30*). If we propagate lineages inheriting either new or old poles forward in time, we obtain an interesting prediction. Provided that λ remains low, a lineage of new pole cells would reach a point where $k_0 = k_1$, and therefore $T_0 = T_1$ (Fig.

1B). The same occurs if we follow a lineage of old pole cells, which reach a point where $k_0 = k_2$ and $T_0 = T_2$. When plotting maternal doubling times against that of each daughter (Fig. 1B), these equilibrium points are represented as the intersection between model predictions and the identity line.

As $\lambda$ increases however, populations reach a threshold where new daughter lineages can still reach the $T_0 = T_1$ equilibrium, but the old daughter lineage can no longer stabilize. As a result, $T_2$ assumed increasingly larger values, until the lineage ceases to divide (Fig. 1C).

**Asymmetry and growth stability of high-persistence mutants**

Because persistence occurs in a small subpopulation of cells, high-persistence mutants are widely used in experimental settings to increase persister frequencies. The most recurrent of these model organisms is the strain *E. coli hipA7*, harboring two mutations on the gene encoding the toxin HipA (*36*). Its use allowed for the first description of a stochastic switch between active and dormant states within microfluidic devices (*2*). However, most studies employing the strain have been performed in batch cultures, offering limited information on the growth physiology of this mutant. As such, we tracked lineages of *E. coli hipA7* growing in the daughter device (Fig. S1) and investigated the effects of asymmetric divisions and stochasticity on the phenotypic heterogeneity of mutant populations.

Overall, the *hipA7* mutation produced no growth anomalies. Populations displayed constant and asymmetric elongation rates over time, with new daughters (0.029 ± 0.002 $min^{-1}$, n = 1,565 cells) elongating faster than old daughters (0.028 ± 0.002 $min^{-1}$, n = 1,505 cells; one-tailed t test, t = 13.37, df = 3,063.1, $p < 0.001$; Fig. S3A). No indication of stochastic switching to a non-growing state was observed under unstressed conditions. Considering pairs of siblings (Fig. S3B), *hipA7* new daughters had shorter doubling times (23.843 ± 1.786 min) than old daughters (24.573 ± 1.749 min; one-tailed paired t test, t = 12.955, df = 1,194, $p < 0.001$). A positive correlation with maternal doubling times was observed for both new (a = 0.443, $p < 0.001$, $R^2 = 0.148$) and old subpopulations (a = 0.349, $p < 0.001$, $R^2 = 0.095$). Since both linear models intersected the identity line, a distinct point of equilibrium for each subpopulation is predicted to exist at these intersections.

To verify whether these lineages displayed physiological stability and long-term proliferation, we analyzed the new and old equilibria. Their stability was determined by estimating the stochasticity ($\sigma_1$) acting on the slope (a) of the linear models represented in Fig. S3B (see Methods and Fig. S2 for details). For $a^2 + \sigma_1^2 < 1$, doubling times of a cell lineage remain stable around the equilibrium over generations, despite the stochasticity present in the system. For *hipA7* populations, both new and old lineages largely satisfied this condition (Fig. S3C), thus displaying continuous proliferation in exponential phase.

Since spontaneous persistence is attributed to stochastic factors, we investigated whether the physiological heterogeneity of *hipA7* mutants harbored more stochasticity than in wild-type bacteria (Fig. S3D to S3F). To estimate the contribution of deterministic and stochastic factors to the heterogeneity of doubling times, we performed the analysis described by Chao et al. 2016 (*43*). This estimate consists of extracting growth parameters describing doubling times through processes of damage inheritance, accumulation, and partitioning. From these parameters, the doubling times of hypothetical symmetric siblings can be obtained for each maternal doubling time. Experimental sibling pairs of new and old daughters are normalized by their symmetric sibling (represented in Fig. S3D and S3E). The separation between new and old distributions is then used to estimate the deterministic component of heterogeneity, whereas the variance of the distributions indicates the stochastic component (see Methods for details). From this analysis, our results indicated a similar level of stochasticity for both wild-type and *hipA7* populations (Fig. S3F). Thus, *hipA7* mutants are not more heterogeneous nor more stochastic than wild-type *E. coli*.

In only one aspect, the high-persistence strain displayed a remarkable distinction from the wild-type. When inoculated into mother machine devices, *hipA7* mutants exhibited a significantly longer time until first division (93.65 ± 26.37 min) than wild-type cells (62.60 ± 14.61 min; one-way ANOVA, $F = 64.112$, $p < 0.001$; Fig. S3G). By the time all wild-type cells had initiated exponential growth, 9.68% of *hipA7* cells remained in lag phase.

This difference in lag duration is crucial for the interpretation of studies in which antibiotic exposure occurs within a few hours of inoculation into fresh medium. To illustrate this point, we inoculated *hipA7* into the mother machine and added 100 μg/ml Ampicillin to the medium immediately after loading (Fig. S3H to S3N). The antibiotic reached the cells before all individuals had entered exponential growth. Growing bacteria were killed, while lag phase bacteria persisted the treatment (Fig. S3M). After 5h of exposure Ampicillin was removed, and former lag phase individuals started growing. While single-cell microscopy experiments allow for an easy differentiation between lag and exponential cells at the onset of the treatment, batch culture experiments require particular care to ensure that stationary cells have not been carried over into the new culture.

#### Damaging treatments failed to produce persisters during exponential phase

Damaging treatments can increase phenotypic heterogeneity and induce differential growth arrest, where old daughters tend to stop replicating earlier than new daughters (*30*). This could provide a deterministic basis for spontaneous persister formation. To test this hypothesis, we tracked cell lineages growing exponentially in mother machine devices and performed a damaging treatment. Photo-oxidative stress was introduced into the system through FITC light pulses (490 nm) applied in 2 min intervals (Fig. 3A). The treatment led to a population-wide elongation rate reduction (two-tailed t test, $t = 11.272$, $df = 1{,}041.5$, $p < 0.001$). As this damage was partitioned over generations, the difference between new and old daughter doubling times increased from 0.897 to 1.365 min ($t = 2.409$, $df = 567.5$, $p = 0.016$). This larger heterogeneity was characterized by a relative increase of deterministic growth predictors, although 66.2% of the variance was purely stochastic (Fig. S4). To verify whether this increase in physiological heterogeneity could aid in persister formation, the damaging treatment was followed by exposure to 100 μg/ml Streptomycin for 6.5 h, and a posterior recovery in unstressed conditions. Throughout the experiment we observed growth arrest (Fig. 3A, detail), but most cells lysed during the recovery period, and no individuals ever resumed growth. Since resuming replication after antibiotic removal is a key feature of persistence, the oxidative treatment failed to produce the phenotype.

We repeated this assay with an alternative combination of damaging and killing agents (Fig. 3B). Bacteria were first exposed to sub-lethal levels of Streptomycin, which induces protein misfolding and asymmetric partitioning between old and new daughters (*23, 37*). Compared to photo-oxidation, this treatment produced higher elongation rate heterogeneity (F-test: $F = 2.319$, $p < 0.001$). Most of this variance was defined by deterministic predictors (77.8%, Fig. S3), mainly the inheritance of maternal damage, which led some individuals to arrest growth (Fig. 3B). Because previous studies have suggested a connection between oxidative damage and persistence to quinolone antibiotics (*15, 19*), we introduced 30 µg/ml Nalidixic Acid as a killing agent. Still, no persisters were observed upon subsequent antibiotic removal. Despite the large physiological variability present in the system (whether mainly deterministic or stochastic), phenotypic heterogeneity alone failed to produce spontaneous persisters during exponential growth.

**SUPPLEMENTARY FIGURES**

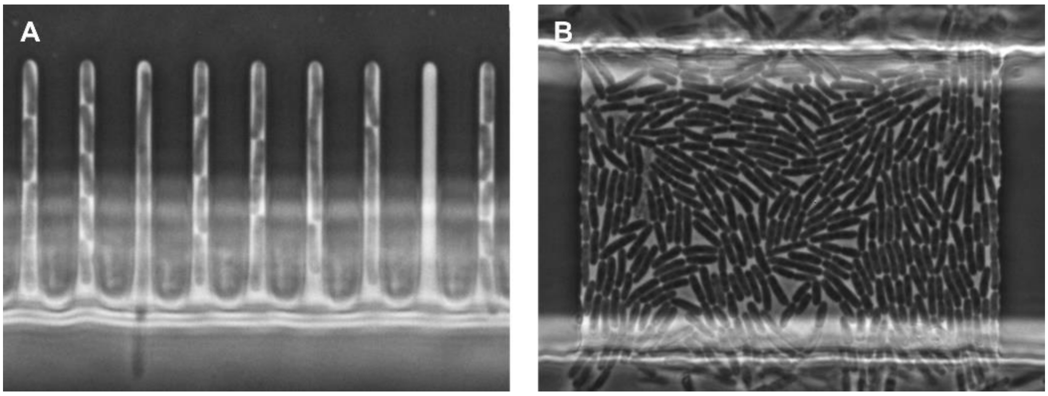


**Fig. S1 Microfluidic devices under phase contrast microscopy. (A)** The mother machine consists of growth wells with one closed end (1.25 × 30 × 1 µm), which trap the old daughter lineage for the duration of the experiments. **(B)** The daughter device consists of large growth chambers (40 × 50 × 0.95 µm) flanked by flow channels, accommodating a population with no bias towards either new or old daughters.

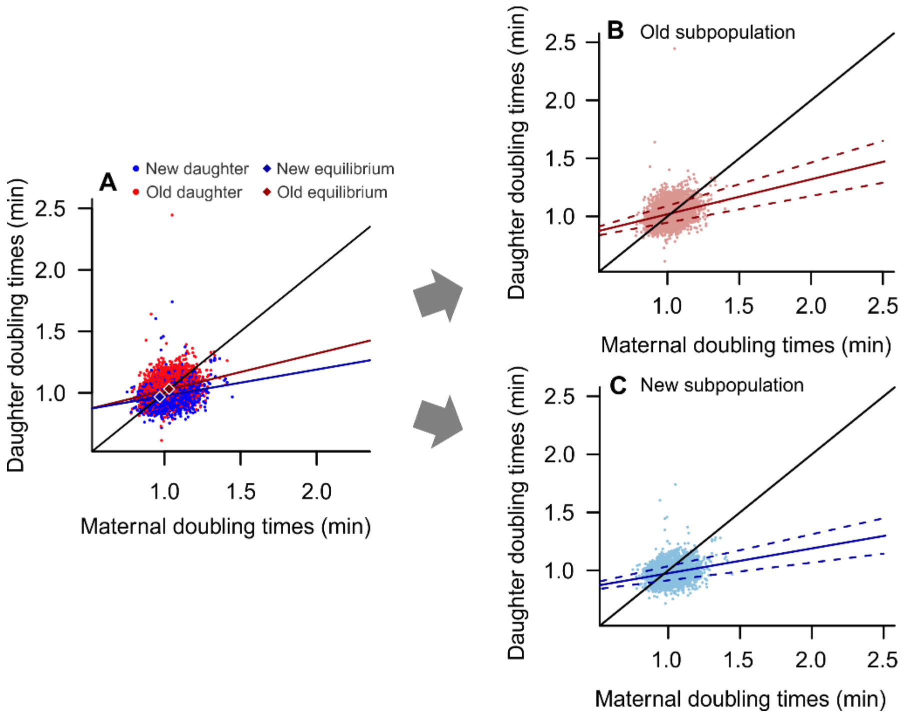


**Fig. S2 Stability of growth equilibria in exponential populations. (A)** Individual data points used in Fig. 2B are presented without binning. The predicted points of equilibrium for new and old subpopulations are indicated through linear regressions between mother and daughter doubling times (see Methods for details). **(B)** Old daughters showed a positive correlation with maternal doubling times ($a = 0.301$, $R^2 = 0.085$, $p < 0.001$; solid red line). The stochasticity present in the system is shown as dashed lines, representing the standard deviation of all slopes between mother and daughters ($\sigma_1 = 0.072$). Because $a^2 + \sigma_1^2 = 0.096$, the condition $a^2 + \sigma_1^2 < 1$ is satisfied and this equilibrium is stable. **(C)** The same pattern was observed for new daughters ($a = 0.214$, $R^2 = 0.062$, $p < 0.001$; solid blue line), with $\sigma_1 = 0.061$. Because $a^2 + \sigma_1^2 = 0.050$, the equilibrium is stable.

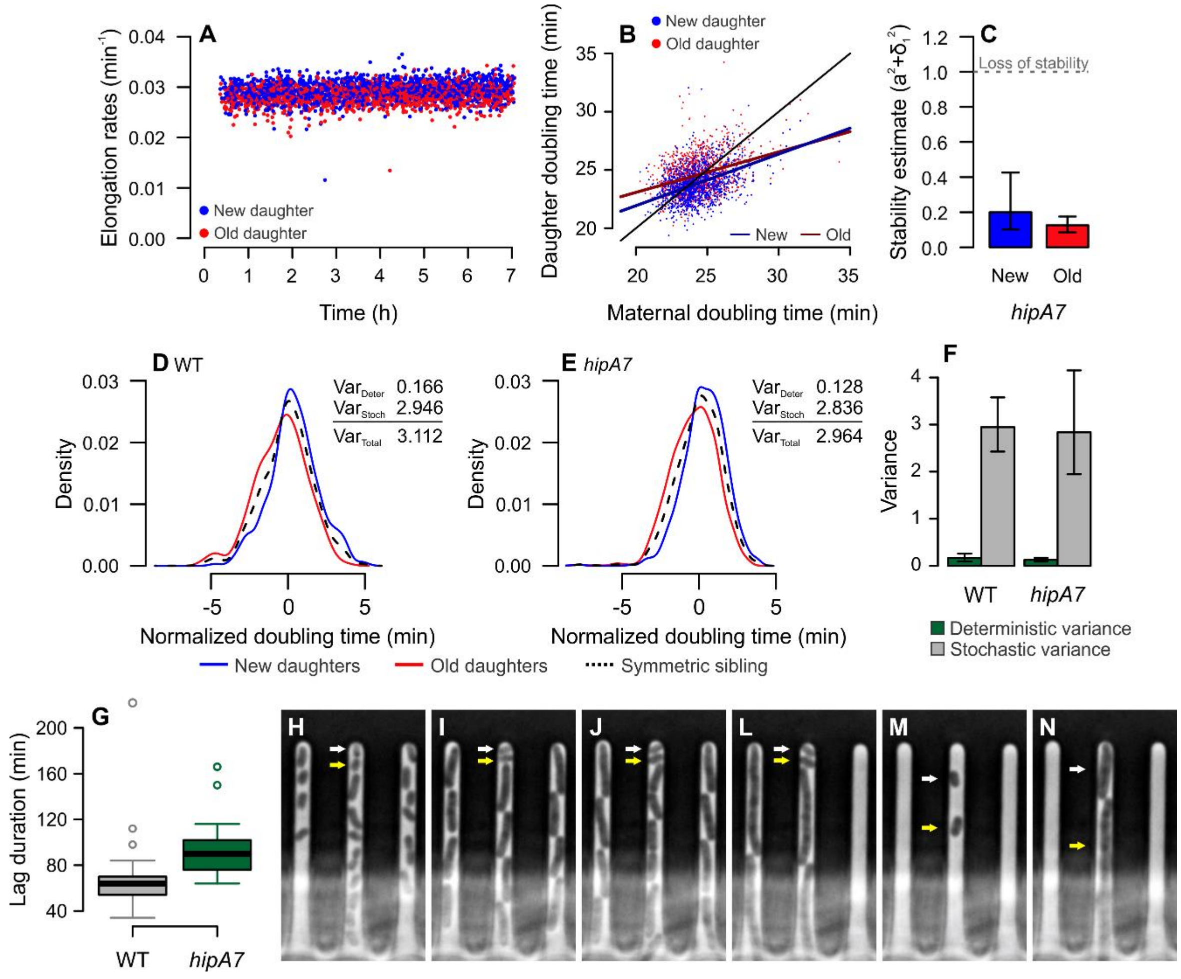


**Fig. S3** ***hipA7*** **exhibits cellular aging and stable growth. A** The high-persistence mutant had stable elongation rates over time, with new daughters displaying faster growth. **(B)** Maternal doubling times showed positive correlation with new and old subpopulations*. Slopes intersecting the identity line suggested points of stable physiology. **(C)** These regressions were used to estimate the stability of equilibrium points according to the slope (a) and stochasticity ($\sigma_1$) for each subpopulation (see Methods). Both new and old lineages had stable equilibria, suggesting continuous growth and replication despite stochastic processes. Error bars = 95% CI. **(D to F)** Partitioning of doubling time variance into deterministic and stochastic components. The distance between distributions provides an estimate of deterministic variance, whereas the mean variance of new and old doubling times distributions estimates the stochastic variance. Compared to wild-type *E. coli*, the *hipA7* mutant did not have higher levels of stochasticity on its growth heterogeneity. Error bars = 95% CI. **(G)** The main physiological distinction between strains was the longer lag phase of *hipA7*. **(H to N)** When cells were exposed to 100 µg/ml Ampicillin shortly after loading, lag phase individuals (arrows) persisted the antibiotic treatment. Time-lapse images are shown in 100 min intervals. The effects of Ampicillin can be observed on (J and L), leading to lysis of growing cells. After 5h, once the antibiotic was removed, persisters started growing (**N**).

* Two points in **B** (daughter doubling times of 51.71 and 60.03 min) are not shown due to plot proportions, but appear as elongation rates in **A**.

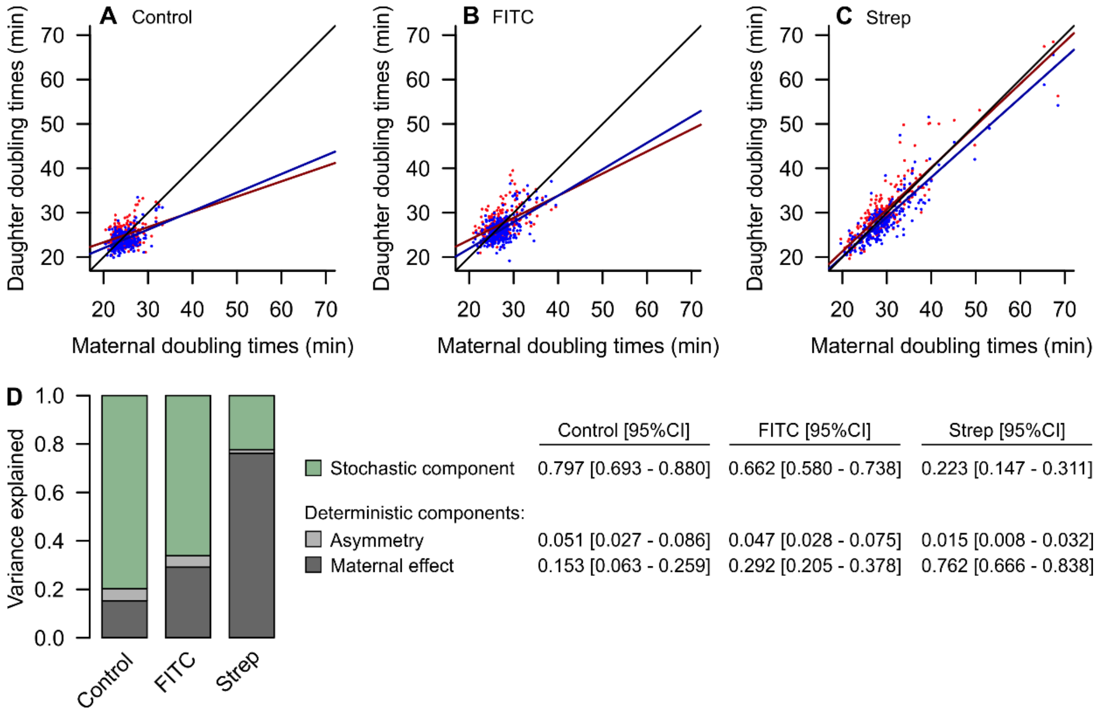


**Fig. S4 Deterministic and stochastic components of the variance for populations under oxidative stress. (A to C)** Phase planes showing the doubling times of new and old daughters as a function of maternal doubling times. **(D)** The total variance of these doubling times was decomposed into its deterministic (asymmetry and maternal effects) and stochastic components through the sums of squared deviations method (see Methods for details). Contrary to the analysis presented in Fig. S3D-F, which compared absolute variance, this estimate considers the relative contribution of each component. Populations exposed to photo-oxidation and Streptomycin showed increasingly more deterministic variability, likely driven by the inheritance of larger damage loads from a mother cell.
